# Mammalian gene expression variability is explained by underlying cell state

**DOI:** 10.1101/626424

**Authors:** Robert Foreman, Roy Wollman

## Abstract

Gene expression variability in mammalian systems plays an important role in physiological and pathophysiological conditions. This variability can come from differential regulation related to cell state (extrinsic) and allele-specific transcriptional bursting (intrinsic). Yet, the relative contribution of these two distinct sources is unknown. Here we exploit the qualitative difference in the patterns of covariance between these two sources to quantify their relative contributions to expression variance in mammalian cells. Using multiplexed error robust RNA fluorescent in situ hybridization (MERFISH) we measured the multivariate gene expression distribution of 150 genes related to Ca^2+^ signaling coupled with the dynamic Ca^2+^ response of live cells to ATP. We show that after controlling for cellular phenotypic states such as size, cell cycle stage, and Ca^2+^ response to ATP, the remaining variability is effectively at the Poisson limit for most genes. These findings demonstrate that the majority of expression variability results from cell state differences and that the contribution of transcriptional bursting is relatively minimal.

## Introduction

Gene expression variability is ubiquitous in all biological systems. In multicellular organisms heterogeneity between different cell types and states confers specialized function giving rise to complexity in whole-system behavior (Eldar and Elowitz, 2010; Raj and van Oudenaarden, 2008; Suo et al., 2018; Symmons and Raj, 2016; Tabula Muris Consortium et al., 2018). Similarly, single-cell organisms and viruses were shown to utilize heterogeneity at the population level to create diverse phenotypes, such as bet-hedging strategies in changing environments (Rouzine et al., 2015; Veening et al., 2008; Vega and Gore, 2014). While variability can provide useful functional heterogeneity in a multicellular organism or cell population, it is not necessarily always beneficial (Raj and van Oudenaarden, 2008; Symmons and Raj, 2016). Unregulated stochastic events, i.e. noise, can limit cells ability to respond accurately to changing environments and can introduce phenotypic variability that can have a negative contribution to overall fitness. Indeed, many biological mechanisms including buffering (Stoeger et al., 2016) and feedback loops (Jangi and Sharp, 2014; Schmiedel et al., 2015) have been suggested to limit the detrimental effect of gene expression variability. Quantification of the different contributions of mechanisms that cause gene expression variability is an important step toward determining to what degree the variability represents uncontrolled “noise” or cellular stratification and function.

Two key contributors of gene expression variability are allele specific sources and global factors related to underlying cell state. Analysis of expression covariance between genes is a powerful approach to decompose gene expression variability into these two classes. Landmark works used this approach to investigate expression variability in bacterial cells, which laid a foundation for decomposing variability into allele-specific (intrinsic) sources and variability that originate from sources that affect multiple alleles and relate to the underlying cell state (extrinsic) (Elowitz, 2002; Paulsson, 2005). This work was later extended to yeast (Raser and O’Shea, 2004) and mammalian systems (Raj et al., 2006; Sigal et al., 2006; Singh et al., 2012). The decomposition into allele-specific and cell state components is not always simple. Allele-specific noise in an upstream component can propagate into downstream genes (Sigal et al., 2006) whereas temporal fluctuations in the shared components can have nontrivial consequences on expression distributions (Paulsson, 2004; Pedraza and van Oudenaarden, 2005; Shahrezaei et al., 2008). Finally, use of the terms “intrinsic” and “extrinsic” is sometimes ill-defined and some models include a “coupled intrinsic” mode as well which is a form of shared variability and hence “extrinsic” (Rodriguez et al., 2019). Despite the sometimes confusing nomenclature, the use of expression covariance to distinguish between allele-specific and shared factors is a powerful decomposition approach.

In addition to covariance based approaches, the relationship between gene expression distribution variance and mean provides a useful quantitative framework to gain insights into sources of expression variability (Munsky et al., 2012). The comparison of expression variability between genes is not straightforward as expression variance scales with its mean. Three statistical tools are commonly used to describe mean normalized variance: the coefficient of variation (CV), coefficient of variation squared (CV^2^), and Fano factor. CV and CV^2^ are both unitless measures where the CV is defined as the standard deviation divided by the mean and the CV^2^ is simply the CV squared, or the variance divided by the mean squared. The CV and CV^2^ are useful to compare the scale of variance between different genes because of their unitless nature. The third measure, the Fano factor, is the variance divided by the mean and therefore not unitless, but it has a special property of being equal to one in the case of a Poisson process. Many biological processes have a variance to mean ratio that is at least Poisson so the Fano factor can define a ‘standard dispersion’, as a result, distributions with Fano factor smaller/bigger than one are considered under/over-dispersed, respectively. Therefore a simple quantification of the distribution variance scaled by its mean can provide key insights into the underlying mechanism generating the observed distribution (Choubey et al., 2015; Hansen, Desai, et al., 2018).

Multiple studies across bacteria, yeast, and mammalian cells measured over-dispersed gene expression distributions. This observation can have two main interpretations. One interpretation is that the observed over-dispersion is simply a result of the superposition of an allele-specific Poisson variability and cell state variability (Battich et al., 2015). The other interpretation is that the allele-specific variability itself is not a simple Poisson process (Corrigan et al., 2016; Dar et al., 2015; Suter et al., 2011; Tantale et al., 2016). The latter interpretation was popularized by the introduction of a simple phenomenological model named the two-state or random telegraph model that represented genes as existing in either “on” or “off” states (Friedman et al., 2006; Fukaya et al., 2016; Kaern et al., 2005; Kepler and Elston, 2001; Lenstra et al., 2016; Molina et al., 2013; Paulsson, 2004; Peccoud and Ycart, 1995; Raj et al., 2006; Sanchez and Golding, 2013; Shahrezaei and Swain, 2008; Suter et al., 2011; Thattai and van Oudenaarden, 2004). More complex models with multiple states were also considered, (Corrigan et al., 2016; Nicolas et al., 2018; Suter et al., 2011; Tantale et al., 2016; Zoller et al., 2015) but the addition of multiple states does not change the model in a qualitative way. These models suggest that transcription should occur in distinct bursts with multiple transcripts generated when the gene is “on”. These two-state models can be described by two overall key parameters: the burst size and frequency that control the resulting gene expression distributions with lower burst frequency and larger burst size contributing to the overdispersion of the underlying distribution. Overall both interpretations, bursting and cell state, can explain the observed over-dispersion and it is currently unclear which one is correct.

The relative scales and sources of variability are very important to understand in the modern world of single-cell highly multiplexed measurements. These new technologies are revealing the complex structure of ‘cell space’ with cells occupying a large array of types (Han et al., 2018; Rosenberg et al., 2018; Tabula Muris Consortium et al., 2018), states (Cheng et al., 2019; Trapnell, 2015), and fronts (Shoval et al., 2012) that reflect functional stratification. Despite our knowledge that cell types and states manifest as gene expression heterogeneity, sometimes total gene expression variability is interpreted as arising from two-state transcriptional bursting alone (Larsson et al., 2019). The gap in our understanding of the relative contribution of cell state and allele-specific factors is hindering progress in assigning functional roles to observed variability (Dueck et al., 2016).

To address this knowledge gap, we utilized the two key properties of expression variability: covariance and dispersion. We measured gene covariance and dispersion using joint measurements of individual cells; where for each cell multiple cell state features were measured as well as a highly multiplexed measurement of gene expression. We used sequential hybridization smFISH (MERFISH implementation) (Moffitt et al., 2016) that allowed us to accurately measure the expression of 150 genes in ~5000 single-cells. Since expression covariance between genes from the same pathway is higher compared to genes that have distinct functions (Sigal et al., 2006; Stewart-Ornstein et al., 2012), we focused on a single signaling network and biological function, Ca^2+^ response to ATP in epithelial cells, a biological response important to wound healing (Funaki et al., 2011; Handly et al., 2015; Handly and Wollman, 2017). The key advantage of Ca^2+^ response is that the overall signaling response can be measured in less than fifteen minutes, a fast timescale that precludes any ATP induced changes in transcription. Using the combined dataset we were able to separate the correlated and uncorrelated components using a simple multiple linear regression model guided by the changes in the covariance matrix. We found that after removing all shared components, the remaining allele-specific variability shows very little over-dispersion for most genes measured. Overall these results indicate that transcriptional bursting is only a minor contributor to the overall observed expression variability.

## Results

To assess the relative contribution of the overall expression variability that stems from allele-specific sources versus underlying cell state variability, we took advantage of the fact that these two sources have different expression covariance signatures. Figure 1 shows simulated data to illustrate how covariance signatures can be utilized to decompose sources of variability. By definition, allele-specific variability is uncorrelated to any other gene whereas variability that is due to heterogeneity in the underlying cell state will likely be shared between genes with similar function (Figure 1A). When transcriptional bursting dominates (Figure 1B top) the shared regulatory factors will have a small contribution, there will be little correlation between genes and the expression variance will remain largely unchanged after conditioning expression level on any cell state factors (Figure 1B top right). The residual intrinsic variance will have a Fano factor greater than one. On the other hand, when cell state variability dominates (Figure 1B bottom), expression between genes will be highly correlated and conditioning the expression on cell state factors will reduce both the variance and correlation between genes. At the limit, when all shared factors are accounted for, the correlation between genes will approach zero and the Fano factor of the residuals will approach one, the Poisson limit (Figure 1B bottom right). When the contribution of bursting and cell state is comparable (Figure 1B middle) conditioning on cell state factors will have some effect but the final Fano factor will be higher than one even when the correlation is zero (Figure 1B middle right). Conditioning on cell state factors has a dual effect on correlation and Fano factor and therefore it is possible to assess whether the conditioning removed all the obvious extrinsic variability. When all the extrinsic variability is conditioned out, one can confidently interpret whether the residual intrinsic variability is under or overdispersed.

**Figure 1.**
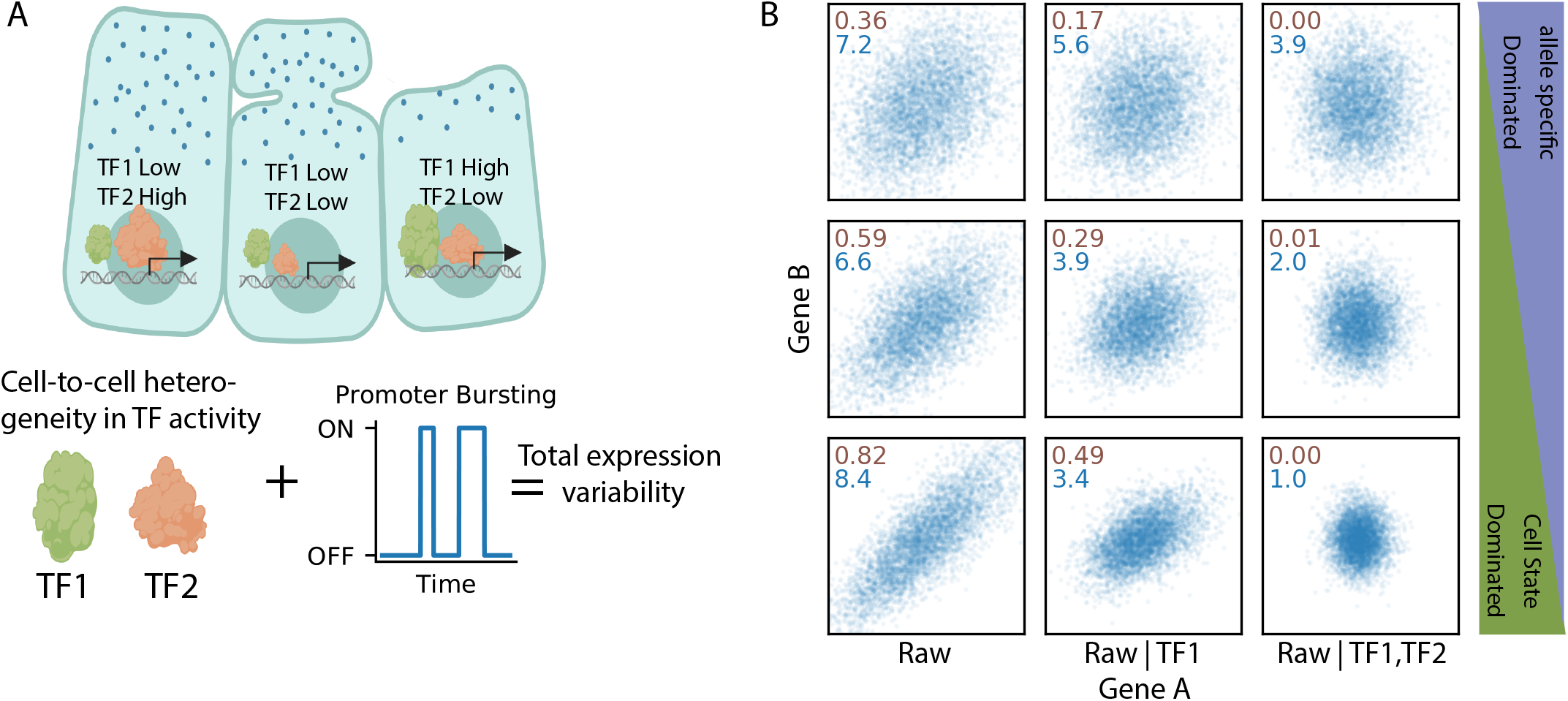
Transcriptional bursting and trans-acting factors are two distinct causes of cell-to-cell heterogeneity. (a) Cartoon depicting that different cells can have different activities of trans-factor (TF) regulatory molecules in addition to the effects of transcriptional bursting. (b) Simulated data showing that variability from shared regulatory factors results in correlation between two genes with three example cases: intrinsic dominated noise (top three panels), mixture of cell-state and allele specific sources (middle three), and cell-state dominated (bottom three). This correlation is diminished when the expression levels are conditioned on the levels of these shared regulatory factors (middle and right). After conditioning on all trans-acting regulatory factors the remaining variability due to transcriptional bursting alone is potentially significantly smaller (right). Inset text is the pearson correlation coefficient between Gene A and Gene B (brown) and the fano factor of Gene A (blue).

To distinguish between the possible situations described above requires accurate highly multiplexed single-cell measurements of gene expression and a sufficient number of cellular features that correlate with the underlying cell state factors controlling gene expression. To achieve this we developed an experimental protocol that combines MERFISH, a multiplexed and error robust protocol of counting RNA transcripts using fluorescent in situ hybridization,(Chen et al., 2015; Moffitt et al., 2016) with rich profiling of the underlying cell state (Figure 2). We used the MCF10A mammary epithelial cell line, which is often used in studies of cellular variability due to their non-transformed nature and their accessibility to imaging (Qu et al., 2015; Selimkhanov et al., 2014). We focused on genes that share biological function: involvement in the Ca^2+^ signaling network, a key pathway important to the cellular response to tissue wounding (Justet et al., 2019; Minns and Trinkaus-Randall, 2016). The two advantages of Ca^2+^ signaling are that 1. we expect that genes that share a function will show a high degree of correlation in their expression levels (Stewart-Ornstein et al., 2012). 2. Ca^2+^ signaling is fast and we can measure the overall emergent phenotype of the network in less than 15 minutes (Figure 2A). In our protocol cells were rapidly fixed after live cell imaging (10-15 min from ATP stimulation to fixation) and therefore the gene expression measured in the same cell is unlikely to have changed as a result of the agonist.

**Figure 2.**
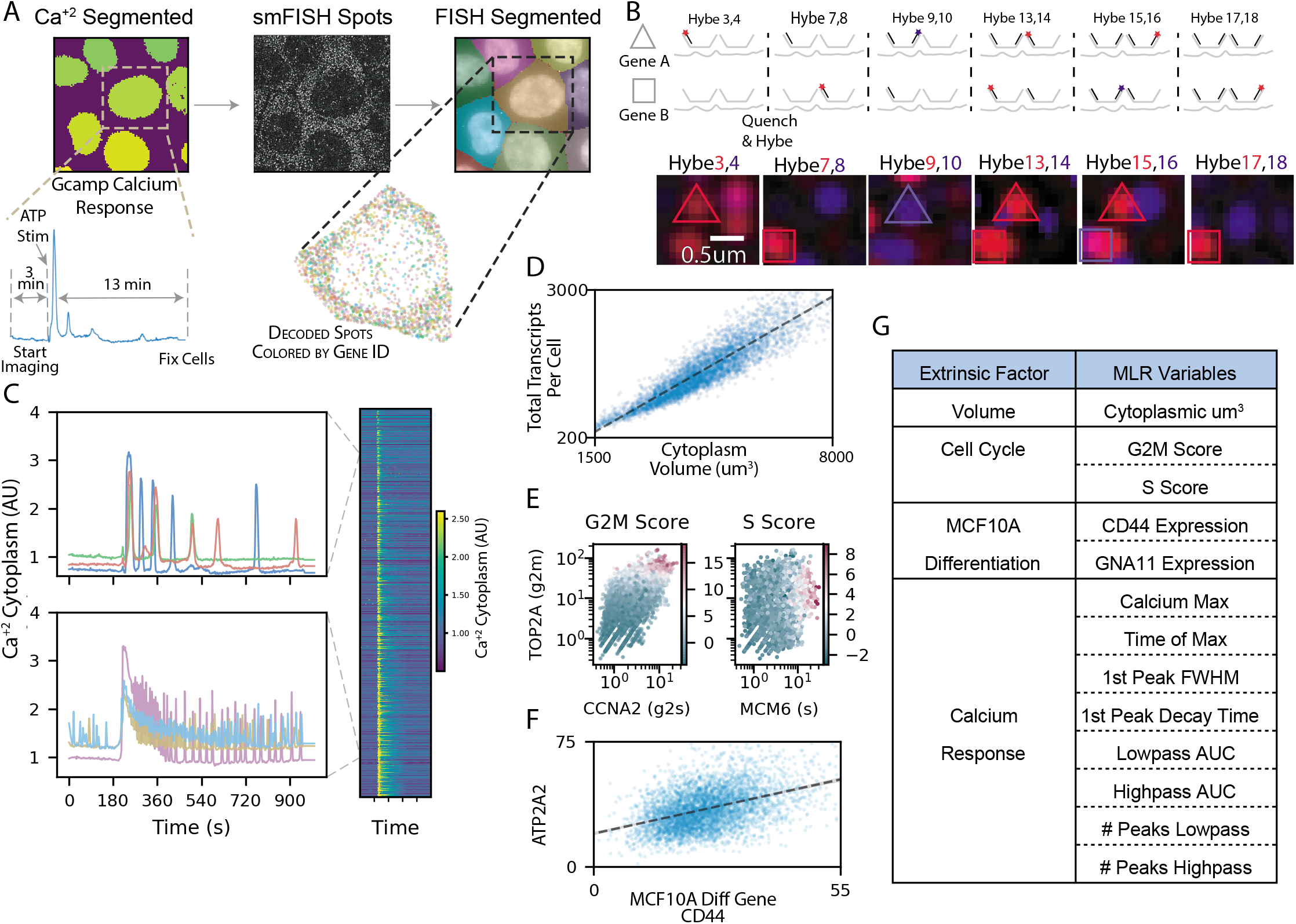
Paired single-cell MERFISH and live-cell calcium imaging. (a) Experimental overview - live cells are imaged for their calcium response to ATP before being fixed and imaged to measure gene expression of 150 genes. (b) smFISH spots are imaged over several rounds of hybridization and aligned such that individual genes are encoded as specific series of dark and bright spots throughout all rounds of hybridization. (c) Left, representative calcium trajectories demonstrating the heterogeneous response to ATP stimulation, top vs bottom left. The Right panel is an image plot of all 5000+ successfully paired to smFISH cells. (d) Cellular volume is measured and the correlation between total transcripts per cell and the cellular volume is shown. (e) Left, shows marker gene expression for cell cycle related genes used to derive a g2m score (coloring). Right, is the same as the left panel with a representative gene used to derive the S score for each cell. (f) Correlation of a representative gene (ATP2A2) with a gene that marks the differentiation status of MCF10A cells (CD44). (g) table of the cell state features categories and the complete list of the 13 factors used in the multiple linear regression (MLR) statistical model.

MERFISH is a multiplexing scheme of smFISH where transcript identity is barcode-based, and the barcodes are imaged over several rounds of hybridization. During each hybridization round, dye-labeled oligos are hybridized to a subset of RNA species being measured, the sample is imaged and RNA appear as diffraction limited spots, then the dye molecules are quenched, and the process is repeated until all barcode ‘bits’ are imaged. By linking diffraction limited spots across imaging rounds, we can decode the RNA barcodes by identifying the subset of images where a bright diffraction limited spot appears at the same XYZ coordinate (Figure 2B). The use of combinatorial labeling allows exponential scaling of the number of genes images with the number of imaging rounds. The scaling is mostly limited by the built-in error correction (Chen et al., 2015). In this experiment, we used 24 imaging rounds (8 hybs x 3 colors) where each RNA molecule was labeled in 4 imaging rounds. An example of the MERFISH data is shown in Figure 2B. Overall we measured the expression of 150 genes including 131 genes annotated as involved in Ca^2+^ signaling network (Bandara et al., 2013; Kanehisa et al., 2019; Kanehisa and Goto, 2000), 17 genes to mark stages of the cell cycle (Whitfield et al., 2002), and two genes that correlate with the sub-differentiated state of MCF10A cells (Qu et al., 2015).

Our decomposition into allele-specific and cell state components is based on conditioning on multiple cell state factors. While, it would be ideal to directly measure the regulatory factors that causatively control gene expression variability, more accessible measurements, e.g. cell size or cell cycle stage, that are correlated with these causative regulatory factors are sufficient for the conditioning process. Given that the genes we probe are related to Ca^2+^ signaling we first extracted key features from time-series of cytoplasmic Ca^2+^ response measured with a calibrated GCaMP5 biosensor (S2 A). The live cell imaging of cytoplasmic Ca^2+^ levels (Figure 2C) showed a highly heterogeneous response, qualitatively and quantitatively similar to previous work on Ca^2+^ signaling in MCF10A cells where we observed a mixed population response with a wide range of response phenotypes (Handly and Wollman, 2017; Yao et al., 2016). We used a feature-based representation of Ca^2+^ response to represent cellular factors that we anticipate correlate with underlying cell state (Figure 2G and S2). In addition to Ca^2+^ features that are specific to Ca^2+^ signaling, we also measured a few global features of the cell that are likely to be correlated with expression changes of most genes. Specifically, we measured cell volume, cell cycle stage, and two markers of MCF10A differentiation status (Figure 2 DEF). As was shown in the past, cell volume strongly correlated with the total number of transcripts per cell (Figure 2D) indicating that at least for some genes cell state factors must be important contributors to their expression variability (Hansen, Desai, et al., 2018; Padovan-Merhar et al., 2015). However, not all genes show the same strength correlation with volume, and some cell cycle genes are more complexly related to volume (S4). Similarly, the cell cycle stage and MCF10A differentiation status were correlated with specific genes (Figure 2EF). Overall we measured 13 different cellular features that will be used to decompose variance in all 131 Ca^2+^ related genes we measured. By focusing on a smaller number of specific features that relate to the Ca^2+^ response augmented by established global cell state features like cell size and cell cycle state we expected to be able to capture most of the expression variability that comes from underlying cell state heterogeneity.

To decompose the observed expression into multiple components we used standard multiple linear regression (MLR) (Battich et al., 2015; Hansen, Desai, et al., 2018). Figure 3A shows the scatter plots of expression of two representative genes (ATP2A2 and RRM1) plotted against cell volume, cell cycle, differentiation markers, and Ca^2+^ feature. The scatter plots show that 1. there is indeed a correlation between expression and some of these cell state features 2. The amount of variance that is explained by each cell state feature can change between genes. Overall the simple MLR model with 13 independent measurements was able to explain between ~15-85% of the observed variance with a median of 0.62 (Figure 3B). To assess the relative contribution of each cell state feature we looked into the relative fraction of explanatory power for each feature category (Figure 3C). Overall, cell volume has the most explanatory power, but for some genes, cell cycle and Ca^2+^ features contribute meaningfully to the explained variance. While some of the features had a small effect in term of the overall variance explained by the feature, in most cases, the effects were very unlikely to be a result of pure random sampling, permutation-based statistical testing showed that most genes measured here are statistically correlated with at least one calcium feature (Figure 3D).

**Figure 3.**
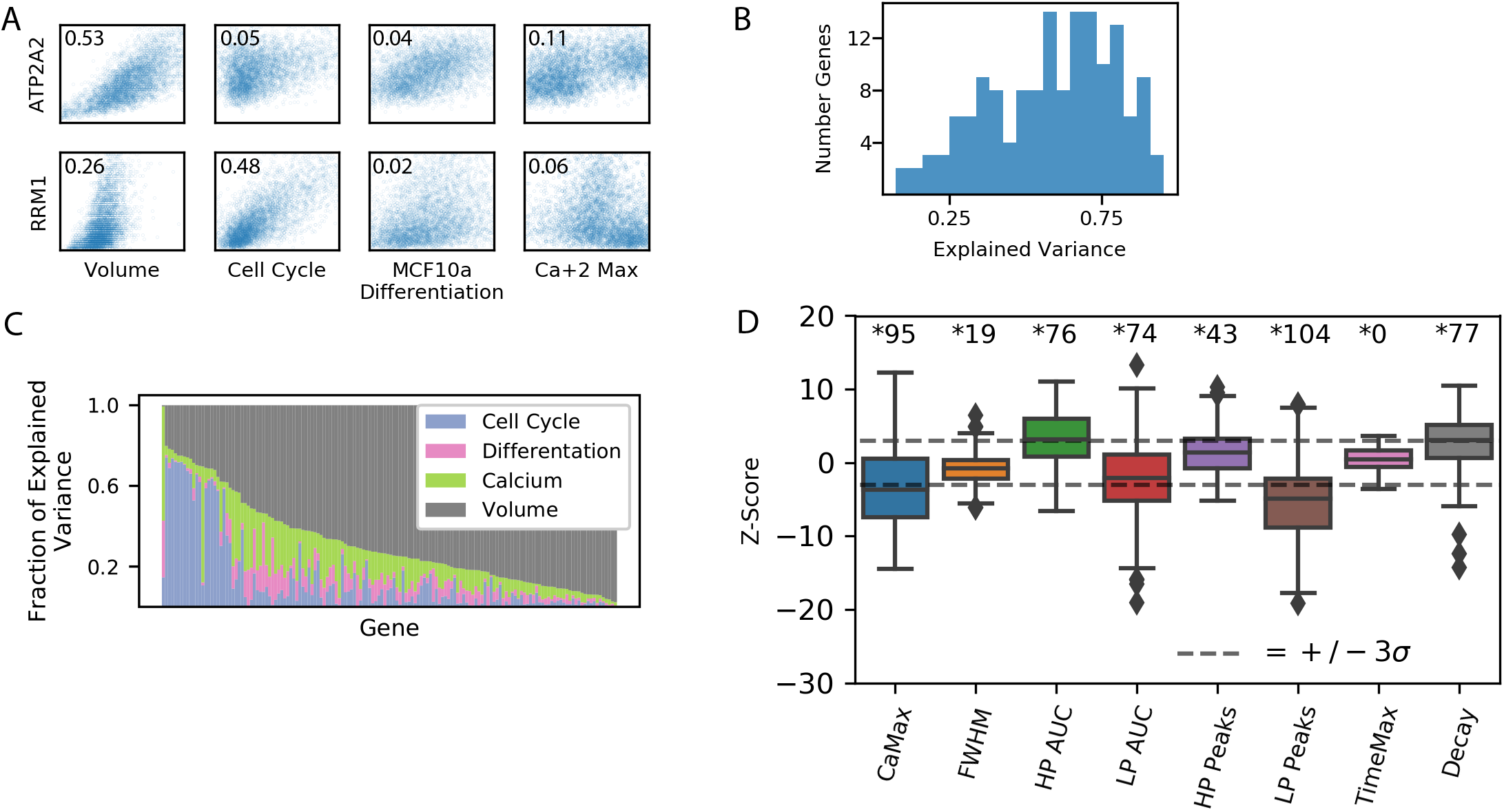
Decomposition of gene expression variability using multiple linear regression. (a) Representative scatter plots of correlation between two individual genes (rows) and different cell state factors (columns). The percent of variance explained by each factor in the MLR model for each gene is annotated in the corner. (b) A histogram of the overall explained variance for each gene. (c) Stacked bar plot showing which cell state features categories contribute to the explained variance of the MLR. (d) The significance of calcium features for each gene were estimated by Z-scoring the slope of the feature in a null distribution of bootstrapped shuffled data slopes. The number of statistically significant genes for each features is shown above (adjusted P-value (Bonferroni) < 0.05).

A key uniqueness of our approach is that gene expression is measured in a multiplexed fashion allowing the estimation of the correlation between genes. Figure 4A shows the correlation matrix of the raw counts, and the counts conditioned on cell state features. As expected, as we increase the number of cell state features included in the MLR, the overall gene to gene correlation goes down. Interestingly, the full MLR model, that only includes 13 identical terms for all genes is able to reduce the overall correlation between gene significantly. To quantify the bulk correlation we measured the amount of variance that is explained by the first two components of a Principal Component Analysis (PCA) (Figure 4B). Without conditioning on any cellular feature, the first two components explain >40% of the variance. This is reduced substantially to <10% of the overall variance, in the full MLR. The substantial reduction in the gene to gene correlation demonstrates that we were able to condition away most of the shared components. Still, the remaining correlation was not completely removed and therefore we added another term to the model that is based on the first two principal components of a PCA analysis after taking all other features into account. These two components most likely represent some cell state features that were not sufficiently captured by our 13 cellular features. With the addition of the last “hidden” feature, the overall variance that is shared is very close to values from shuffled data. Overall the analysis of expression covariance demonstrates that our simple MLR sufficiently captures most of the information related to cell state that is required for conditioning expression distribution.

**Figure 4.**
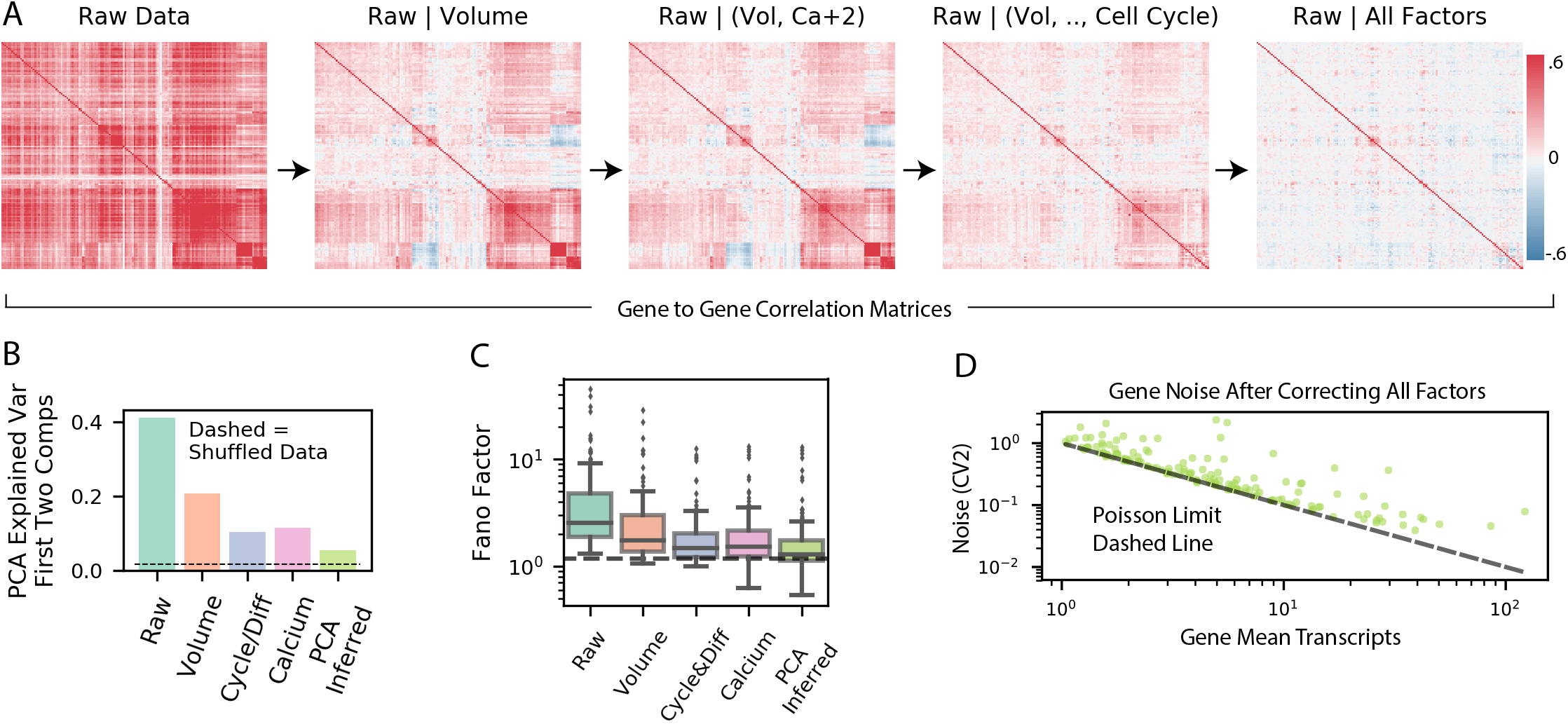
Residual variability from MLR models contains significantly less covariation between genes and close to poisson variability within individual genes. (a) Gene-gene correlation matrices showing the reduction of covariance after conditioning on cell state features. (b) Explained variance of first 2 components of PCA for each stage of MLR models showing reduction in shared variability with increasing number of cell state factors. (c) Fano factor distributions at different levels of cell state conditioning are shown as boxplots. Dashed line is the Poisson expectation. (d) Scatter plot of residual gene expression coefficient variation squared for each gene after decomposition of all cell state features. Poisson expectation is shown as dashed line.

Finally, we wanted to determine the overall dispersion remaining in the allele-specific gene expression distribution. The allele-specific variability is estimated as the residual variability in the raw gene expression counts after conditioning on cell state factors. As we increase the number of cell state features we conditioned on, we saw a substantial reduction in the distributions of Fano factor magnitudes (Figure 4C). When all 13 cell state features and the two hidden features estimated based on PCA are included, the Fano factor is very close to one for most of the genes. Note that we do not perform any correction for technical noise so the limit of one is only theoretical. Similarly, analysis of the coefficient of variation square (CV^2^) vs the expression means on a log-log plot shows that all genes are very close to the Poisson limit (Figure 4D). The proximity to the Poisson limit is similar across all expression levels. Therefore, these data indicate that super-Poissonian transcriptional bursting plays a very minor role in allele-specific variability. It is unclear if the few genes that do show over-dispersion whether they have significant levels of transcriptional bursting or whether our conditioning procedure failed to sufficiently remove cell state effect.

## Discussion

Here we analyzed the relative contribution of gene specific variability that arises from transcriptional bursting, i.e. episodic synthesis of multiple transcripts from a gene, and variability that is shared among multiple genes. Our approach is enabled by very rich single cell measurement that include live cell Ca^2+^ response to ATP, global cell state factors such as size and cell cycle stage, and the expression level of 150 genes all in the same single cells. Using this data, we were able to decompose gene expression variability into gene-specific and and cell state components. We show that after removing covariability from gene expression distributions, the remaining variability follows a simple Poisson model. The residual allele specific variability is not over-dispersed and therefore not consistent with models of transcriptional bursting where a gene is actively transcribed only during a small fraction of time.

The popularity of the transcriptional bursting model is evident by the large number of papers that fit the entire RNA and protein distributions to the two state model without considering other sources of variability (Dey et al., 2015; Molina et al., 2013; Skupsky et al., 2010; Suter et al., 2011). In other cases, cell state was considered using dual reporters (Sigal et al., 2006; Strebinger et al., 2018), assuming timescale separation (Dar et al., 2012), or conditioning on forward scatter (Sherman et al., 2015). However, without multiplexed expression measurements it is difficult to determine whether conditioning on cell state was done to completion. The high goodness of fit of the two-state model to uncorrected or partially corrected distributions that shows substantial bursting could simply be a case of over interpretation of model fit. RNA binding systems, such as MS2, allow direct live-cell observation of transcription bursting, and many groups have observed burst-like punctuated transcription (Corrigan et al., 2016; Ferguson and Larson, 2013; Fritzsch et al., 2018; Muramoto et al., 2010). While direct visualization is compelling, it is unclear if punctuated transcriptional events are due to stochastic transition of promoter state, as suggested by two state model, or due to stochasticity in the activity of an upstream regulatory element. Furthermore, difficulty in quantifying the number of mRNAs synthesized in each such event make it difficult to distinguish between a two-state model and a one state model with a low rate of transcription that will generate a Poisson distribution. In fact, our results are consistent with recent measurements that showed that TTF1 mRNA is generated in “bursts” of 1-2 mRNA (Rodriguez et al., 2019). Furthermore, the two alleles of TTF1 showed coordination between these bursts suggesting that the observed transcriptional events are coupled through trans-regulatory factors. Finally, temporal changes in global rates of transcriptions (Shah et al., 2018; Skinner et al., 2016) can also make the interpretation of a single allele temporal reporter challenging. It is important to note that our work focuses on genes that encode for calcium signaling activity and might not represent all genes, such as reporters controlled by viral promoters (Dar et al., 2012; Singh et al., 2010) and genes that are key to cellular differentiation (Hansen and van Oudenaarden, 2013; Ochiai et al., 2014). Overall it is advisable to use more caution when interpreting gene expression variability as evidence of transcriptional bursting.

Our measurements are based on cytoplasmic RNA and it is possible that mechanisms related to RNA processing reduce the dispersion of RNA distribution in the cytoplasm after it was generated in an over-dispersed manner through bursting (Battich et al., 2015). Cells include a large number of RNA binding proteins many with unknown function and it is possible that some function as part of post-transcriptional noise reduction mechanisms (Hansen, Wen, et al., 2018). However, some of the proposed mechanisms such as nuclear export of RNA were shown to act as amplifiers of observed dispersion (Hansen, Desai, et al., 2018). Therefore the degree by which post-transcriptional mechanism can be used to reduced expression noise is an important open question. Until additional data will help clarify the ubiquity of such mechanisms, the most parsimonious interpretation is simply that RNA synthesis does not happen in large allele-specific bursts.

Recent technological advances in the ability to measure single cell gene expression with scRNAseq and sequential smFISH approaches are providing an unparalleled view into the underlying “cell state space”. The distribution of cells in “cell state space” and the definition of cell types and states within this space are key open research areas that will likely to further grow in importance with further improvements in single cell measurement technologies (Eng et al., 2019; Wagner et al., 2016). Our work has two important implications on our understanding of this “cell state space”, at least with regards to the heterogeneity of a single cell type: 1. All the shared variability was reduced using only a simple representation of cell state as 13 linear coefficients. Furthermore, most of these 13 features had only a very small contribution to the overall explanatory power suggesting that cell state distribution can be represented by few latent dimensions. An observation that emboldens efforts to learn the cell state manifold (Moon et al., 2018). 2. Expression noise, i.e. unregulated variability in gene expression that is a result of stochastic biochemical interactions in effect defines a “resolution limit” of the cell state space. Our results indicate that the highly heterogeneous distribution of cells within cell state space is likely not due to the inability of cells to control their expression levels rather our work indicates functional stratification of cells within this space. Collectively these contributions pave the way to a more rigorous definition of cell state that is based on concepts of signal to noise where the signal is represented by regulated differences between cells and noise is due to unregulated stochastic events. Such definitions will help identify the functional role of cellular heterogeneity.

## Acknowledgements

We are thankful to Jeffrey Moffitt for his extensive feedback about the MERFISH method, and Anna Pilko for making the mCherry-Gcamp construct and cell line. This work was funded by NIH grants to RW EY024960 and GM111404.

## Author Contributions

RF and RW conceptualized the experiments and data analysis. RF performed the experiments and performed data analysis. RF and RW wrote and edited the paper.

## Methods

### Contact for Reagent and Resource Sharing

Further information and requests for resources and reagents should be directed to and will be fulfilled by the Lead Contact, Roy Wollman.

### Experimental Model and Subject Details

The MCF10a cells used in this study are Homo Sapien, female cells with the RRID: CRL-10317. This cell line has not been authenticated, but bought directly from ATCC. Cells were grown in complete media: DMEM/F12 media (Gibco) supplemented with 5% Horse Serum (Life Technologies), EGF 20ng/mL, hydrocortisone 0.5ug/mL, cholera toxin 0.1ug/mL, insulin 10ug/mL, and Penicillin/Step 100U/mL referred to as complete media.

### Cell Culture

MCF10a cells were grown in complete media (above) and passaged at 70-90% confluency. Cells were seeded onto coated 40mm #1.5 coverslips (Bioptech) and grown to confluence in 5mm diameter PDMS wells before changing media to complete media without EGF and 1% horse serum, instead of normal 5%, 6-8 hours before imaging. Coating solution consists of sterile filtered 10ug/mL fibronectin, 10ug/mL bovine serum albumin, and 30ug/mL type I collagen in DMEM.

### mCherry GCamp5 Fusion Construct Creation

For pPB - mCherry vector construction a PCR product encoding GCaMP5 sensor incorporating the CaMP3 mutation T302L R303P D380Y and no stop codon (Addgene plasmid #31788) was directionally ligated into pENTR/D-TOPO vector (Invitrogen K243520) resulting in pEntry_GCaMP5G construct. (For:caccATGGGTTCTCATCATCATCATCATCATGGTATGGCTAGCATGAC, REV: TTACTTCGCTGTCATCATTTGTACAAACTCTTCGTAG) pEntry_GCaMP5G was linearized with PCR reaction using standard Phusion^®^ Hot Start Flex 2X Master Mix (NEB Cat# M0536L) protocol (FOR: cgcgccgacccag, REV: ctcgagggatccggatcctcccttcgctgtcatcatttgtacaaac). PCR product was then subjected to DpnI digestion (NEB cat# R0176S) and gel purification with Zymoclean Gel DNA Recovery Kit (ZYMO cat#D4001). A sequence encoding mCherry and a5’ linker was PCR amplified (FOR : gaggatccggatccctcgagAccatggtgagcaagggc REV: aagaaagctgggtcggcgcgcttgtacagctcgtccatg). mCherry2-C1 was a gift from Michael Davidson (Addgene plasmid # 54563). GeneArt Seamless Cloning and Assembly Enzyme Mix (Invitrogen cat# A14606) was used to assemble a construct encoding for GCaMP5 sensor fused with a short linker to mCherry called pENTRY-GCaMP5fusedmCherry. LR recombination between this entry clone and a custom gateway PiggyBack transposon vector with 1 μl LR Clonase II enzyme (Invitrogen: cat #11791020) resulted in the final construct of pPB_CAG_GCaMP5fusedmCherry_blast.

### mCherry GCamp5 Fusion MCF10A Cell Line Creation

To generate stable cell lines constitutively expressing cGamp5fusion-mcherry,

MCF10A cells grown in the standard conditions and co-transfected using Neon transfection system (Invitrogen cat#MPK1025) and transposase expression vector pCMV-hyPBase (Sanger institute) in the 4:1 ratio with 0.625 ug of transposase and 2ug of transposon plasmid per well in 6 well dish. Electroporation parameters:

Pulse voltage (v) 1,100 2003

Pulse width (ms) 20

Pulse number 2

Cell density (cells/ml) 2 x 10^5

Transfection efficiency 45%

Viability 65%

Tip type 10 μ

Stable, polyclonal cell populations were established after blasticidin selection (10 μg/mL).

### Coverslip Modification

40mm coverslips (Bioptech) were allyl-silane functionalized according to (Moffitt et al., 2016) which briefly consists of washing coverslips in 50% methanol and 50% 12M HCl, and then incubating at room temperature in 0.1% (vol/vol) triethylamine (Millipore), 0.2% (vol/vol) allyltrichlorosilane (Sigma) in chloroform for 30 minutes. Washing with chloroform then 100% ethanol and air drying with nitrogen gas. These were stored in a desiccator for less than a month until use.

### Calcium Imaging

Cells were stained with 0.1 ug/mL Hoescht for 20 minutes then rinsed with imaging media. Each well was imaged and stimulated consecutively as follows: image 3 minutes of Gcamp before stimulating with 6uM ATP in imaging media then imaged for another 13 minutes. Gcamp was imaged every 2-3 seconds and Hoechst was imaged every 4 minutes for segmentation. Immediately following imaging of a well, that well was fixed with 4% formaldhyde in PBS. The next well was imaged, and then the previously imaged/fixed well was washed 3X with PBS.

### Sequential FISH Staining

PDMS wells were removed and cells were briefly fixed for 2 minutes, washed 3X with PBS, and then permeabilized with 0.5% Triton X-100 in PBS for 15 minutes. Coverslips were washed 3X with 50mM Tris and 300mM NaCl (TBS), and then immersed in 30% formamide in TBS (MW) for 5 minutes to equilibrate, all the liquid was aspirated from the petri dishes, and 30uL of 75uM encoding probes and 1uM locked poly-T oligos were added on top of the coverslip and a piece of parafilm was place on top of the coverslip to evenly spread the small volume over the surface and prevent evaporation. The entire petridish was also sealed with parafilm and incubated at 37C for 36-48 hours. The parafilm was removed and the coverslip was washed 2X with MW buffer with 30 minute incubation at 47C for both washes. A 4% polyacrylamide hydrogel was then cast to embed the cells before clearing with 2% SDS, 0.5% Triton X-100, and 8U/mL proteinase k (NEB P8107S), according to previously published methods. Coverslips were incubated in clearing buffer for 24 hours then washed 3X in TBS for 15 minutes each at room temperature. (Moffitt et al., 2016)

### Sequential FISH Imaging

smFISH staining was imaged on a custom modified Zeiss Axiobserver Z1 body with Andor Zyla 4.2 sCMOS camera and 1.4NA 63 Plan-Apo oil immersion objective. Illumination light was provided by luxeon rebel LEDs (Deep Red, Lime, Blue, and Royal Blue) to excite Cy5, Atto565, Alexa 488, Hoechts, and 200nm Deep Blue fiducial markers. The microscope was controlled by micro-manager (Ausubel et al., 2001) and custom MATLAB software. Automated washing during sequential rounds of hybridization was accomplished by using a previous published setup (Moffitt et al., 2016; Moffitt and Zhuang, 2016). Briefly, FC2 bioptech flow chambers were attached to a gilson minipuls peristaltic pump pulling liquid from reservoirs attached to hamilton MVP valves. The pump and valves were controlled with arduino, and serial commands with Python https://github.com/ZhuangLab/storm-control/tree/master/storm_control/fluidics. This setup was used to automatically wash cells with TBS, then 2mL of TCEP (Sigma) in TBS incubated for 15 minutes, then rinse with TBS, then flow in 2mL of wash buffer (10% ethylene carbonate in TBS with 2mM Vanadyl Ribonucleoside Complex (NEB)), followed by 3mL 3nM readout probes in wash buffer incubated for 15 minutes, then rinsed with 2mL wash buffer, then 1mL of TBS, and finally 3mL of imaging buffer. Imaging buffer is 0.15U/mL rPCO (OYCO), 2mM PCA (Sigma), 2mM Trolox (Sigma), 50mM pH 8.0 Tris-HCl, 300mM NaCl, and 40U/mL murine rnase inhibitor (NEB).

### FISH Oligo Pool Design Amplification

Oligopools were ordered from CustomArray. The oligos were designed using previously published software (Moffitt and Zhuang, 2016). Briefly, design involves selecting 30bp regions with 40-60% GC for each target gene that maximizes specificity of the oligo by finding shared 15-mer substrings against all other transcripts in the human genome. These regions are concatenated with sequences for 3 readout probe binding sequences and flanking 20bp primers.

Probes were amplified according to the another previously published work (Wang et al., 2018). Briefly, limited cycle qPCR with a T7 promoter on the reverse primer. The PCR was terminated 1 cycle after saturation during the extension phase. PCR product was column purified, then *in vitro* transcription further amplified the oligos (NEB Quick High Yield Kit), t7 reactions were purified with desalting columns, and converted to ssDNA with Maxima RT H- (Thermo).

### Gcamp Image Processing

Cell nuclei were segmented using custom Python 3.6 scripts. Cell nuclei were segmented using the Hoechts staining. Nuclear images were low pass filters with gaussian of sigma 5 pixels. Then regional maxima were found with corner_peaks from scikit-image these peaks were used as seeds in a watershed of the negative intensity of the images, and thresholded with otsu of the smoothed nuclear images. This was repeated for each time point and the centroid of each nuclear mask was tracked across time using linear assignment. Segmented nuclei were used as masks to calculate the mean intensity within each cell mask in the Gcamp channel and also the channel for mCherry-fusion expression marker for Gcamp. Finally Gcamp values were divided by the mCherry values to give expression normalized calcium trajectories.

### Calcium Trajectory Feature Extraction

Calcium trajectories were processed with wavelets to find lowpass, smoothed, and highpass trajectories by thresholding coefficients of different scale wavelets. Peaks were detected in the lowpass and highpass with scipy’s find_peaks and prominence thresholds of 0.1 and 0.15 respectively. Decay time of the first major peak after ATP stim was calculated, FWHM of the first peak after ATP was calculated, the AUC of highpass and lowpass was calculated with numpy’s trapz, the maximum of each calcium was calculated, and the time of maximum was also calculated from smooth trajectories.

### Alignment to Live Cell Images

EM microgrids (G400F1-Cu EMS) were glued (23005 biotium) to 40mm Bioptech coverslips. These grids were imaged in brightfield to determine the stage coordinate of fiduciary marks on the microgrids. A rotation and translation transformation was fitted between the live cell and smFISH coordinates of microgrid fiduciary marks. This ensured that we imaged the same FOVs, but additional alignment was performed after imaging. smFISH images were downscaled until they had a pixel size matching the live cell imaging (63x vs 10x with same Andor Zyla Camera so 6.3X downscaling). Cross-correlation template matching with live cell templates and smFISH candidate images was performed iteratively with range of rotational angles (−5 to +5 degrees) in order a second set of ‘image’ translations and rotations that maximize the cross correlation scores. A threshold was then applied and downsampled images were stitched together and overlaid to confirm successful alignment.

### smFISH Image Alignment

All rounds of hybridization contained 200nm blue beads (F8805 ThermoFisher) that were imaged in addition to smFISH oligos.

First the coordinates of putative beads were determined with subpixel accuracy by upsampling images by a factor of 5 (~20.5 nm pixel size) and finding peak coordinates of normalized cross-correlation between a gaussian ‘bead template’ and bead images in 3D. Next a translational transformation was estimated from these putative beads with a custom algorithm designed to be robust to false detection of beads. Briefly, neighborhoods of beads with a radius of maximum shift (100 pixels), were found and the differences each of these pairs was calculated. Next, the bead coordinate differences were density clustered and bead pairs from the largest cluster were used in a least sq error optimization of translation vector that minimizes residual of all bead pairs after translation. This fit was performed in 3D and any FOVs with a residual >0.5 pixels XY or 1.2um (3 frames) in Z were discarded.

### Chromatic Aberration Correction

Tetraspeck (4-color) 100nm beads were imaged in all channels used for smFISH imaging. The subpixel centers of these beads were found as described above, and the misalignment of channels was calculated as a function of the XY image coordinate. Images were then interpolated in 2D to correct for systematic differences between channels. (Mostly only necessary at edges of the images due to large camera sensor size).

### Gene Calling

Spots were called with a reimplemented algorithm deeply inspired by (Moffitt et al., 2016), and code is available at https://github.com/wollmanlab/PySpots. Images were taken every 0.4um in Z, but groups of 3 images 1 above and below the current Z slice being processed were maximum projected to form a pseudo Z slice to be further processed. Then two Z slice were skipped before form another pseudo Z slice. This local max projections help gene calling perhaps do to making the imaging more robust to misaligned images, or uncorrected planarity issues in the objective. Second, fiduciary 200nm beads were used to fit XYZ translation transformations described in the image alignment section, and all psuedo Z slices were warped to correct for chromatic aberration and translations from stage reproducibility error. Registered and chromatic aberration fixed images were then high pass filtered by subtracting a gaussian convolution with sigma 2.2 pixels from the original images. These high pass filtered images were then deconvolved for 20 iterations of lucy richardson deconvolution using the flowdec package.(Czech et al., 2018) Finally after deconvolution the images were blurred by gaussian convolution with a sigma of 0.9 pixels. The output at this step for each site imaged is a matrix of (2048, 2048, 24, #Z) elements. Where 2048 is the image width and height, 24 is the number of codebits used to encode gene identity (3 colors X 8 rounds sequential hybridization) and #Z is the number of pseudo Z slices. Next, each Z slice was processed separately on a per pixel basis to assign each pixel as its gene identity or as background. This process was done by dividing each of the 24 images by the 95th percentile of that image to make the intensities for different codebits more similar, and L-2 normalizing each pixel. Then for each pixel the euclidean distance to L-2 normalized codebit vectors was calculated, and if that distance was less than the volume of a nonoverlapping hypersphere for all codewords (0.5176) then the pixel was classified as that closest codeword. This approach is essentially testing whether the intensities from all 24 codebits point in the direction of a particular codeword in 24-dimensional space. Finally, these classified images (2048, 2048, #Z) were segmented to collect groups of connected components with that same gene label. Finally genes calls were thresholded on the number of pixels for each group of connected components, and the average intensity of the set of connected components.

### Calculation of Cell Volume

A 3-D histogram of gene calls for each cell was calculated and smoothed with a gaussian filter of 10 pixels. The number of voxels (1um, 1um, 1um) with at least 0.5 RNA was calculated and used as the volume for each cell.

#### Simulation of Gene Variance Decomposition

For each of the three combinations of cell state and allele-specific noise simulations there were three transcription factors and two genes simulated. Transcription factors were poisson distributed, and genes were simulated as gamma distributions with shapes dependent on additive combinations of transcriptions 1, 2, and 3. The scale of the gamma distributions were varied to control the amount of ‘allele specific variability’, and the amount of gene correlation was controlled by the fraction of shape shared between genes.

For each of the 3 combinations of different noises there were 4 linear models fitted using python statsmodels ols package. For each gene a model was fitted for gene ~ tf1 and gene ~ tf1+tf2. Then the residuals from the fit were adjusted by adding back the mean of expression for that gene, and these mean adjusted residuals are the distribution of the gene conditioned on tf1 or (tf1, tf2).

#### Cell Cycle Features

Cell cycle features were calculated using the scanpy package (Wolf et al., 2018).

#### Gene Variance Decomposition

The same method (linear model residuals) as in the simulation was used to decompose variance for gene expression. In order to investigate residual correlations between genes with different sets of conditioning variables, the decomposition was repeated from different combinations of feature combinations. The first stage involved only gene ~ volume, and then gene ~ volume + s_phase + g2m_phase…finally for the inferred features we used PCA components #1 and #2 as features: gene ~ pca_comp1 + pca_comp2.

#### Statistical Test of Calcium Feature Significance

Volume adjusted gene expression counts were fitted with a linear model based on calcium features. For every gene separate and every calcium feature separately a shuffled linear model was also calculated. That is, for each calcium feature and gene many bootstrap models were estimated where a single calcium feature was shuffled and the model was fitted. The slopes of these fitted models on shuffled data formed a null distribution, and then the p-value of the feature for that gene was considered (100-Qtile(unshuffled slope in shuffled bootstraps)) where 0 is 1/#Bootstraps.

---

**Supplementary Figure 1.**
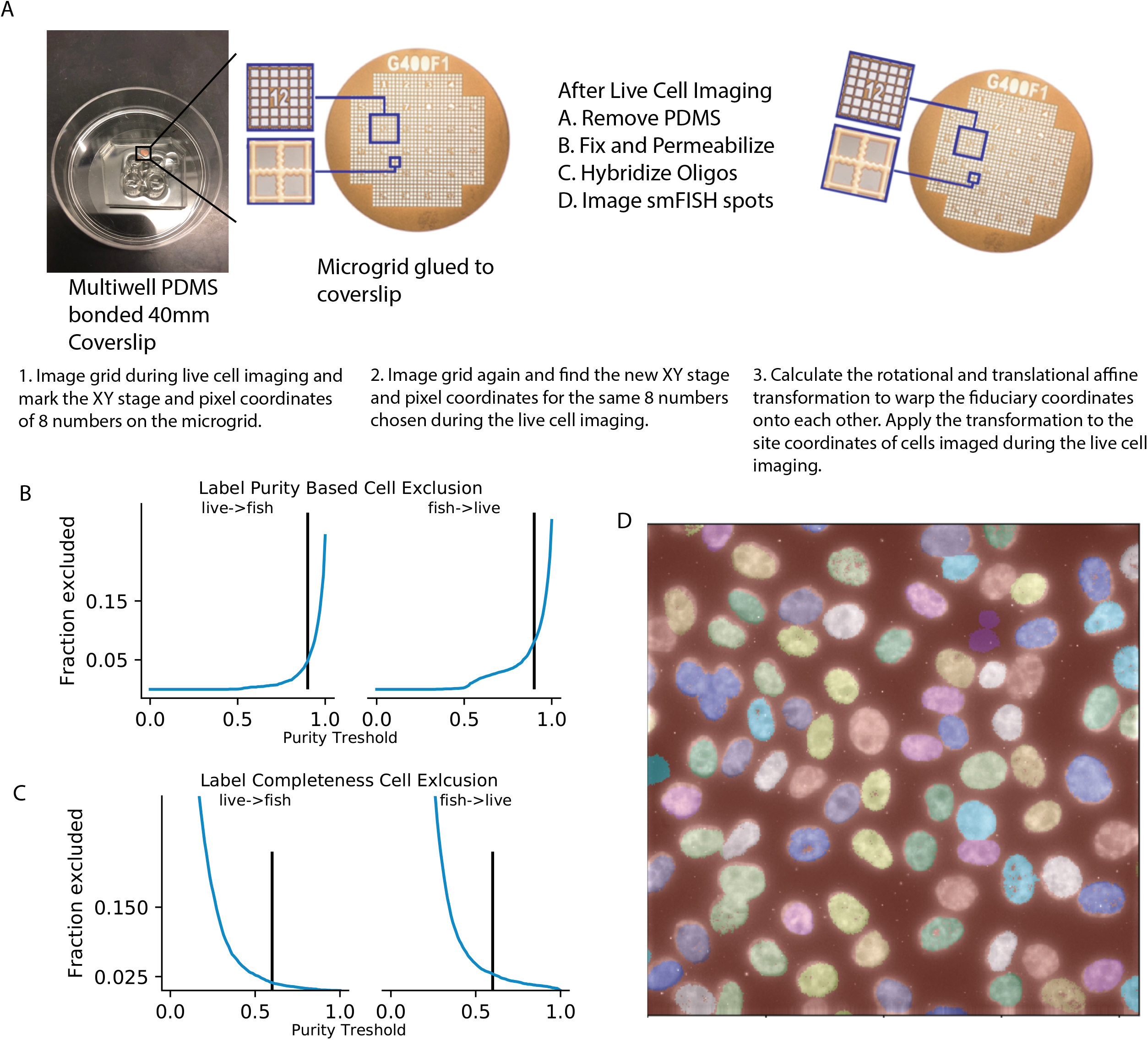
Alignment of calcium images cells with smFISH imaging cells and quality control filtering of cells. (a) PDMS wells and the fiduciary grid attached to a 40mm coverslip (left). Example of orientation of grid during the live cell imaging (middle). Right panel shows that during the smFISH imaging the grid can be rotated, but coordinates of the fiduciary numbers are identifiable. (Below) description of the steps taken to align the fiduciary grids. (b) Purity of label is the fraction of pixels in the calcium nucleus labels that are the same as the nucleus labels from the FISH imaging (left), and vice-versa (right). Vertical lines represent thresholds used to discard cells which were not uniquely mapped between live calcium and fixed FISH imaging. (c) Label completeness is the fraction pixels that were non-zero in the paired segmentation and vice-versa (left, right). Vertical lines were the thresholds of completeness used in quality filtering of alignments.

**Supplementary Figure 2.**
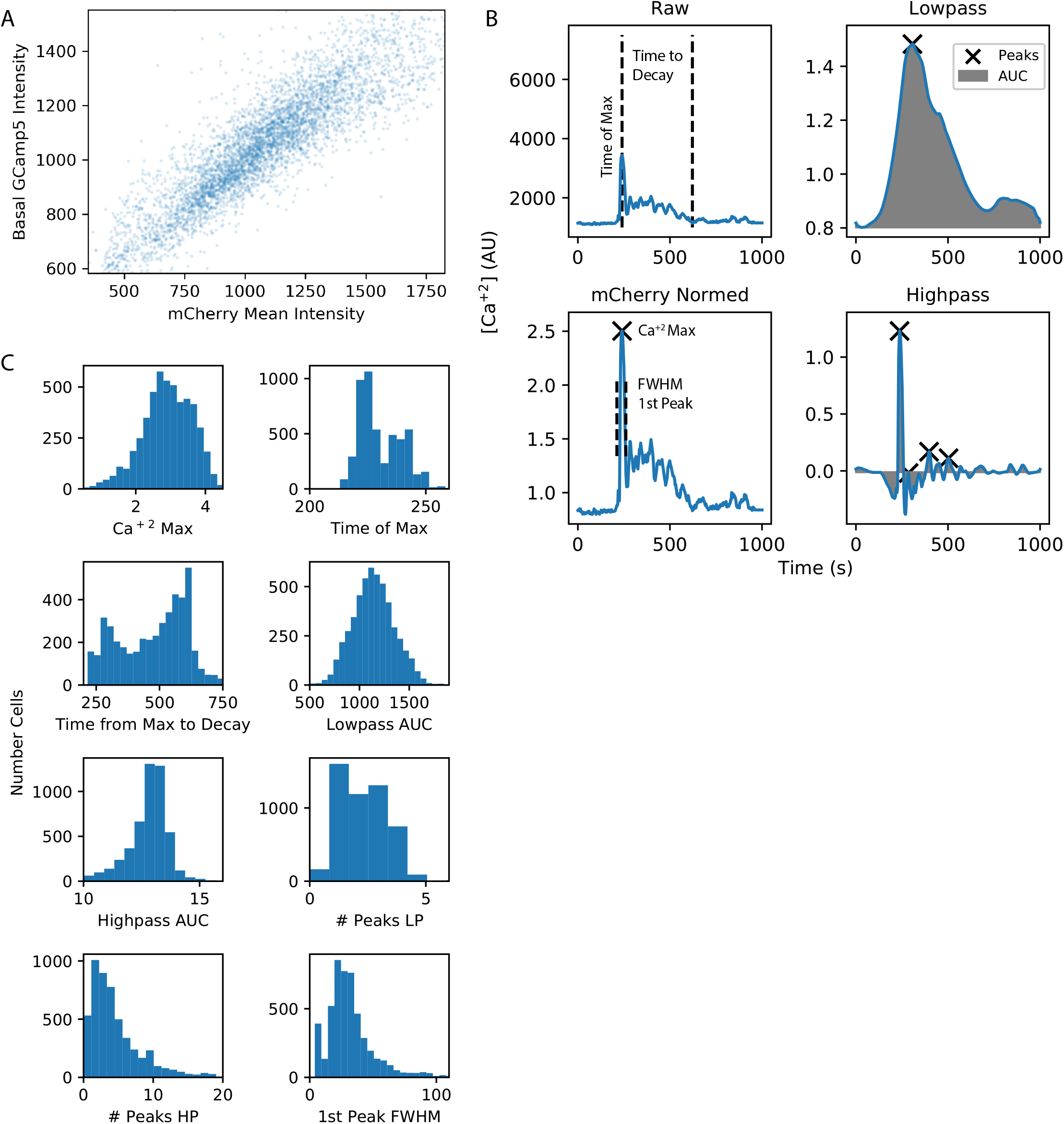
Feature based representation of calcium trajectories. (a) Ca^+2^ is measured with GCaMP sensor fused with mCherry. The high correlation between basal GCaMP intensity and mCherry intensity indicates that mCherry intensity can be used to normalize sensor expression variability. (b) Top left shows raw GCaMP Ca+^2^ trajectories with the Time to Decay of 1st peak calcium feature, and the Time of Max feature for the example trajectory (dotted lines). Bottom left shows the mCherry normalized trajectory with the FWHM of 1st peak and the intensity value of Ca^+2^ Max feature for an example trajectory. Top right shows the lowpass filtered trajectory and the AUC (grey) as well as where the peaks in the lowpass are (x). Bottom right shows the same AUC (grey) and number of peaks (x) but for the high pass filtered trajectory. (c) Histograms of each calcium feature for all ~5000 cells are shown in each subpanel.

**Supplementary Figure 3.**
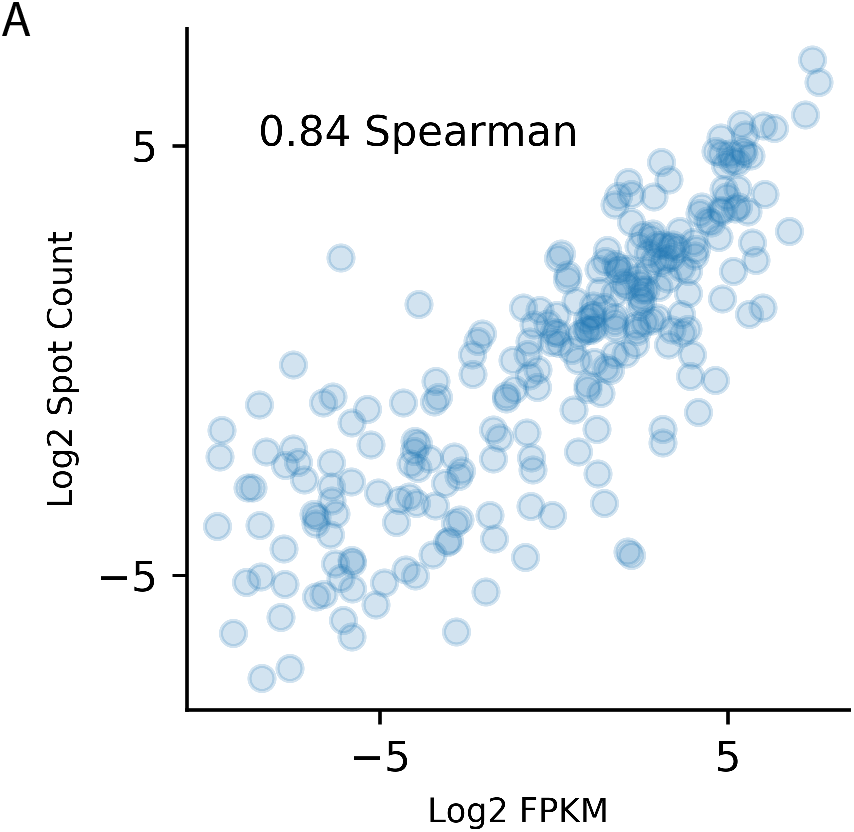
Sequential hybridization smFISH is accurate measure of gene expression. (a) The scatter plot and correlation of RNA-Seq FPKM vs spot counts from the sequential smFISH measurements.

**Supplementary Figure 4.**
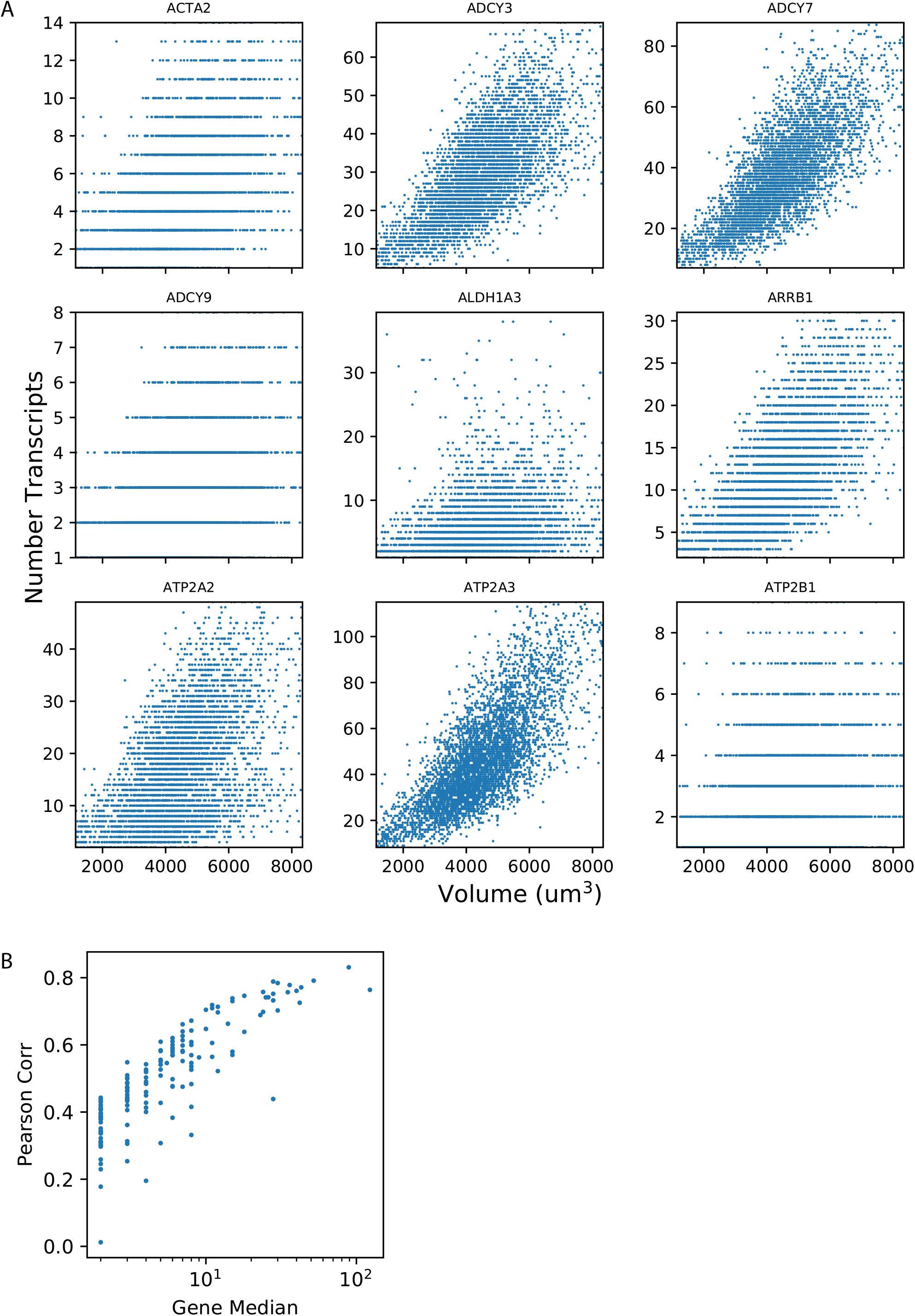
The relationship between volume and gene expression counts for different genes. (a) Nine randomly selected genes are shown as a scatter plot of volume vs spot counts per cell. (b) The Pearson correlation of volume and gene expression as a function of the gene’s median expression.

